# CytoSet: Predicting clinical outcomes via set-modeling of cytometry data

**DOI:** 10.1101/2021.04.13.439702

**Authors:** Haidong Yi, Natalie Stanley

## Abstract

Single-cell flow and mass cytometry technologies are being increasingly applied in clinical settings, as they enable the simultaneous measurement of multiple proteins across millions of cells within a multi-patient cohort. In this work, we introduce CytoSet, a deep learning model that can directly predict a patient’s clinical outcome from a collection of cells obtained through a blood or tissue sample. Unlike previous work, CytoSet explicitly models the cells profiled in each patient sample as a set, allowing for the use of recently developed permutation invariant architectures. We show that CytoSet achieves state-of-the-art classification performance across a variety of flow and mass cytometry benchmark datasets. The strong classification performance is further complemented by demonstrated robustness to the number of sub-sampled cells per patient and the depth of model, enabling CytoSet to scale adequately to hundreds of patient samples. The strong performance achieved by the set-based architectures used in CytoSet suggests that clinical cytometry data can be appropriately interpreted and studied as sets. The code is publicly available at https://github.com/CompCy-lab/cytoset.

## 1 Introduction

High-throughput single-cell technologies, such as flow and mass cytometry, have shown particular promise in numerous translational applications in systems immunology (Davis et al., 2017), such as, pregnancy (Aghaeepour et al., 2017), aging (Alpert et al., 2019), and recovery from surgical trauma (Ganio et al., 2020). In clinical settings, these technologies can simultaneously measure between 20-40 markers at single-cell resolution (Spitzer & Nolan, 2016), across multiple patient samples. The protein measurements collected for each cell facilitate the phenotyping and functional characterization of diverse immune cell-types. Such information can be further used to predict a patient’s clinical outcome, or classification (e.g. healthy or sick).

Based on the financial and time expenses of modern single-cell assays, it is increasingly important to develop robust machine learning methods that enable clinically-meaningful predictions from a set of profiled cells. Such tasks therefore defines a computational challenge of linking the set of multiple single-cell measurements per patient to their respective clinical outcomes. To address this, two major classes of techniques have been proposed. The so-called ‘gating-based’ methods (Bruggner et al., 2014; Lun et al., 2017; Weber et al., 2019; Stanley et al., 2020) first cluster cells across all patient samples into homogeneous subsets according to the expression of the measured proteins. From the resulting clusters, a simple feature vector is typically engineered for each sample, reflecting the relative distribution (or frequency) of cells across each cluster. Such engineered feature vectors can therefore be used for downstream tasks, such as regression (Aghaeepour et al., 2017) and classification (Stanley et al., 2020; Bruggner et al., 2014; Ganio et al., 2020).

The feature vector defined in this ‘gating’ step has two critical properties: (1) it has a user-specified length, despite the fact that the number of cells across different samples is often quite variable; (2) the order of cells in each sample does not affect the corresponding feature vector. Although manual or automatic gating generates features that can be input to a standard machine learning model (e.g. logistic regression), these approaches also present several problems. First, unsupervised clustering requires specifying the number of clusters in advance and most methods do not scale well to millions of cells. Second, clustering often suffers from high variance by different random seeds (Stanley et al., 2020). Finally, the gating step builds on the assumption that all of the useful and clinically-relevant information can be represented by computing the distribution of cells across clusters in each sample. In summary, the ‘gating’ step imposed through clustering introduces a strong inductive bias in the feature extraction step.

To address these limitations, two gating-free methods (Arvaniti & Claassen, 2017; Hu et al., 2019) were recently proposed. Unlike the clustering, or primarily ‘gating’-based approaches, these techniques operate on the single-cell level and try to aggregate information across all single cells in a learnable way. For example, CellCNN (Arvaniti & Claassen, 2017) uses convolutional neural networks (CNNs) as an end-to-end model to learn the associated phenotype from multi-cell input. Specifically, CellCNN uses a 1d-convolution layer to project the measurements of each cell to an embedding space and then applies a pooling layer to aggregate information across multiple different cells. CytoDx (Hu et al., 2019) uses a two-level linear model to predict clinical outcome across individual cells. CytoDx operates on each individual cell and generates the predictor for the sample. At the sample level, CytoDx uses logistic regression to link the predictor and the corresponding clinical outcome.

Although these techniques can successfully identify and achieve strong classification accuracy in small datasets, they often do not adapt well to larger datasets, which are likely to contain bias from batch effects and noisy measurements. In this paper, we propose a new deep learning model called CytoSet to predict clinical outcomes from single-cell flow and mass cytometry data. CytoSet can robustly predict clinical outcomes in four flow and mass cytometry datasets, even with only a limited subset of cells (on the order of thousands per sample). The contributions of CytoSet can be summarized as follows:

1. CytoSet is an end-to-end model that can predict clinical outcomes directly on a set of cells profiled in each patient sample. The prediction task with cells as inputs, contrasts the gating-based approaches, which train a model based on the engineered feature vectors.
2. CytoSet uses the permutation invariant network architecture inspired by the seminal Deep Sets work introduced in Ref. (Zaheer et al., 2017) to extract information from set-structured cytometry data.
3. CytoSet achieves superior classification performance on four flow and mass cytometry datasets, in comparison to two baseline methods (CellCNN and CytoDX).

## 2 Methods

In this section, we introduce the CytoSet model. We begin by introducing the notation used throughout the paper. Next, we describe the intuition behind set modeling and and explain why it is useful for classification tasks on single-cell flow and mass cytometry data. We then provide a comprehensive description of CytoSet, including the network architecture and the loss function. Finally, we compare the CytoSet model to two existing methods, CellCNN (Arvaniti & Claassen, 2017) and CytoDx (Hu et al., 2019).

### 2.1 Notation

We let ***X***, **X**, and ***x*** denote a set, a matrix, and a column vector, respectively. We further use *x* or *X* to represent scalars. Given a matrix **X**, we use **X**(*i*, :) and **X**(:, *j*) to represent the *i*th row and *j*th column of **X**, respectively. The (*i, j*)th element in **X** is denoted by **X**(*i, j*). Given a vector ***x***, we use ***x***(*i*) to denote the *i*th element of ***x***. We use **X**^⊤^ and ***x***^⊤^ to represent the transpose of matrix **X** and vector ***x***, respectively.

### 2.2 Set Modeling

For set-structured data, each sample is a collection of unordered data points denoted by 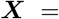 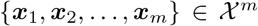, where *m* is the cardinality of the set ***X*** and 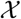 denotes the domain of each element ***x***_*i*_. For example, a set of multiple dimensional points, such as *d*-dimensional single-cell measurements can be viewed as a set. Since different sets are equal only if they have the same elements, models that map a set to some target values must preserve the following permutation invariant property:

#### Definition 1 (Permutation Invariant)

*Let 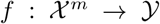 be a function, then f is permutation invariant iff for any permutation π*(·), *f* (***X***) = *f* (*π*(***X***)).

In addition, we also introduce another kind of mapping that is important for building permutation invariant functions in set modeling:

#### Definition 2 (Permutation Equivalent)

*Let 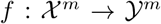 be a function, then f is permutation equivalent iff for any permutation π*(·), *f* (*π*(***X***)) = *π*(*f* (***X***)).

One natural way to encourage permutation invariance is to cluster the elements of a set and output some summary statistics as the feature vector, such as, the proportion of cells assigned to each cluster. Although clustering can preserve permutation invariance, it cannot be trained in an end-to-end fashion using back-propagation because the operation in clustering is non-differentiable. To solve this problem, Refs. Zaheer et al. (2017) and Edwards & Storkey (2017) propose permutation invariant neural network architectures by incorporating permutation equivalent layers and set pooling layers. The set-pooling layers play a key role in preserving permutation invariance and in aggregating information over the elements of a set. Additionally, Ref. Zaheer et al. (2017) also showed the general form of functions that have permutation invariant properties. Recently, Ref. Lee et al. (2019) proposed a new transformer-based architecture that uses attention mechanisms for both encoding and aggregating features in a set.

### 2.3 Problem Formulation

Let ***X*** = {***x***_1_,…, ***x***_*m*_} denote a collection of multiple cells profiled with a high-throughput single-cell technology, such as flow or mass cytometry. Here, *m* is the number of cells and each ***x***_*i*_ ∈ ℝ^*p*^ represents the vector of expression measurements for each of the *p* proteins in cell *i*. In flow and mass cytometry experiments, the order in which cells are profiled has no biological relevance, so such a collection of cells, ***X***, can be viewed as a set rather than as a data matrix. Given a training cytometry dataset with *N* example cells 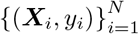 where ***X***_*i*_ is the *i*-th sample that has *m_i_* profiled single cells and *y_i_* ∈ {0, 1} denotes the clinical outcome of ***X***_*i*_, our objective is to learn a permutation invariant function *f*_Θ_(·) that can assign the corresponding label *y_i_* to each set instance, ***X***_*i*_.

### 2.4 Network architecture

Here, we introduce our CytoSet model. The architecture is illustrated in Figure 1. Like CellCNN, CytoSet also takes a set of cells from each individual sample as the input. Let **X** denote the multi-cell input matrix of *k* measured cells, which is randomly drawn with replacement across all of the individual samples ***X***, i.e. **X** ⊆ *X*. This sampling step will generate multiple instances of each sample in the model’s training. This is necessary because (1) the number of cells typically varies across the input samples; (2) the collective number of cells across the original input samples is too large to fit into a GPU. In addition, the diversity across sampled subsets further increases the number of cells involved in training and help the model scale to larger datasets (Amores, 2013). Our experimental results demonstrate that our CytoSet model is robust to the total number of sampled cells, *k* (see section 3.3).

**Figure 1:**
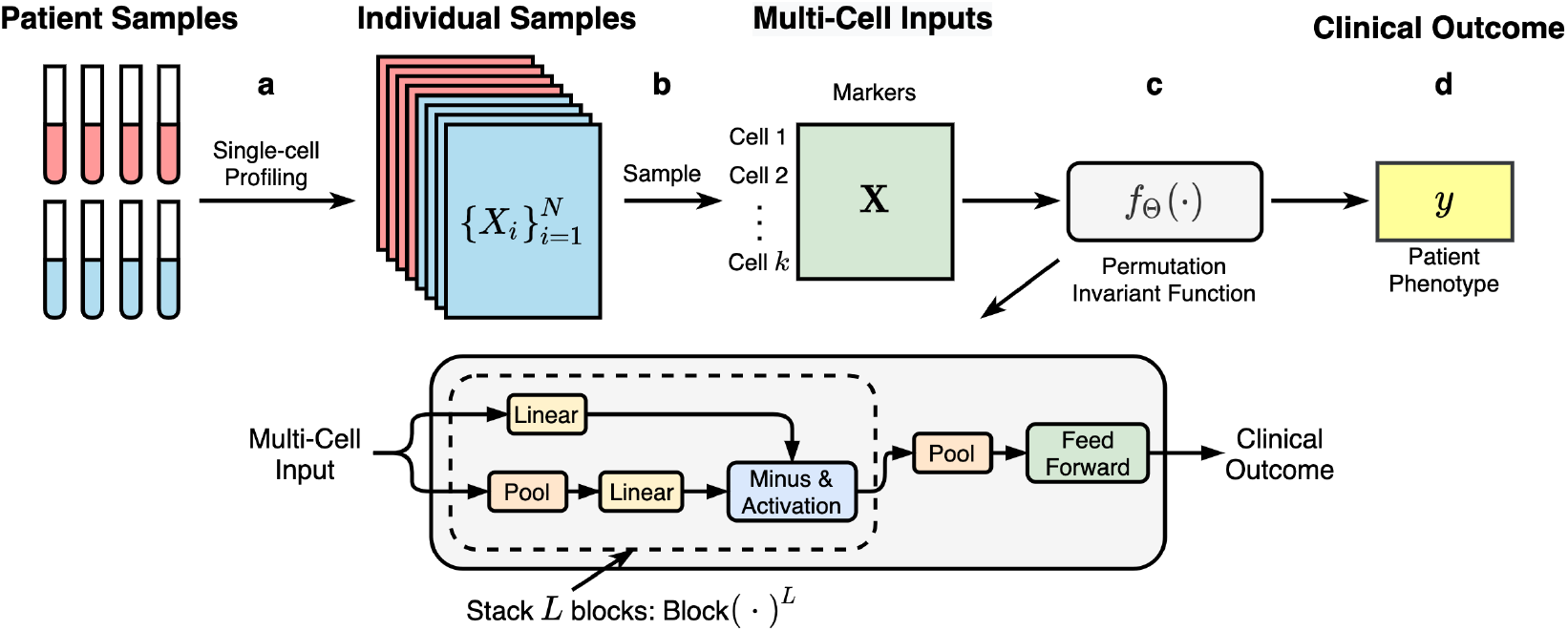
An illustration of the model architecture of CytoSet. (**a**) The collected patient samples are profiled with multiple proteins (markers) at a single-cell level. (**b**) A subsample of all profiled cells across samples is used for training the model. Each multi-cell input is sampled with replacement from an individual sample and has a fixed number of cells, *k*. (**c**) The details of the permutation invariant network architecture used in the CytoSet model. (**d**) The learned feature vectors of multi-cell inputs are used to classify the clinical outcome of patients (e.g. patient phenotype).

Inspired by Ref. Zaheer et al. (2017), the network *f*_Θ_(·) starts by stacking several permutation equivalent blocks that transform the representation of each element in the set. We denote the transformation given by each block in the network as a function, Block(·). This set-based transformation layer will update the set representation iteratively with *L* blocks, and the output by the *l*th block can be denoted as,

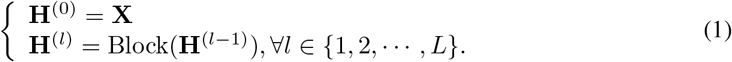

Here, **H**^(*l*−1)^ is the output of the (*l* − 1)th block. In each block, we also add a residual connection to stabilize the network training. After *L* permutation equivalent blocks, a pooling operation is then performed within the rows of **H**^*L*^ ∈ ℝ^*k×d*^, and yields a fixed-length embedding vector for the set ***X***. We denote the embedding vector as ***v*** ∈ ℝ^*d*^, defined as,

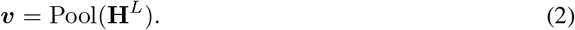

In the implementation of (2), we can either use max or mean pooling. Max pooling returns the maximum value of each column in **H**^*L*^ as,

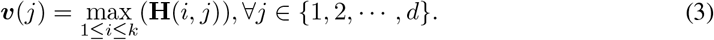

Alternatively, mean pooling returns the mean vale of each column in **H**^*L*^ as,

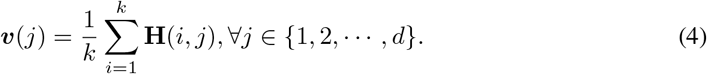

As was previously described in CellCNN (Arvaniti & Claassen, 2017), max pooling can measure the presence of cells with high response, while mean pooling can approximate the frequency of particular cell subsets. In our experiments, we consistently used max pooling to aggregate information among cells. Finally, the embedding vector ***v*** is connected to a feed forward network (fully connected layers) to predict the clinical outcome (e.g. the binary phenotype), *y*.

### 2.5 Loss Function

After several set transformation layers followed by a pooling layer, each set can be represented by a single vector. We use multilayer perceptron neural networks as the classification layers in our model. Letting ***h*** denote the input to the final classification layer in *f*_Θ_(·), we can write the conditional log likelihood of the label *y_i_*, given the set ***X***_*i*_ and model parameters Θ as,

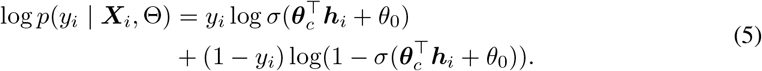

Here, ***θ***_*c*_ and *θ*_0_ are the classification parameters in Θ and *σ*(*z*) = 1/(1 + exp(−*z*)) is the *sigmoid* function. The presented permutation invariant deep neural network is trained by minimizing the following binary cross-entropy loss function,

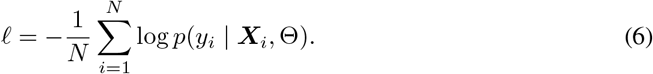

We have thus far introduced all of the components used in our CytoSet model (Figure 1). The details of the training algorithm used for CytoSet is given in Algorithm 1.

**Algorithm 1.**
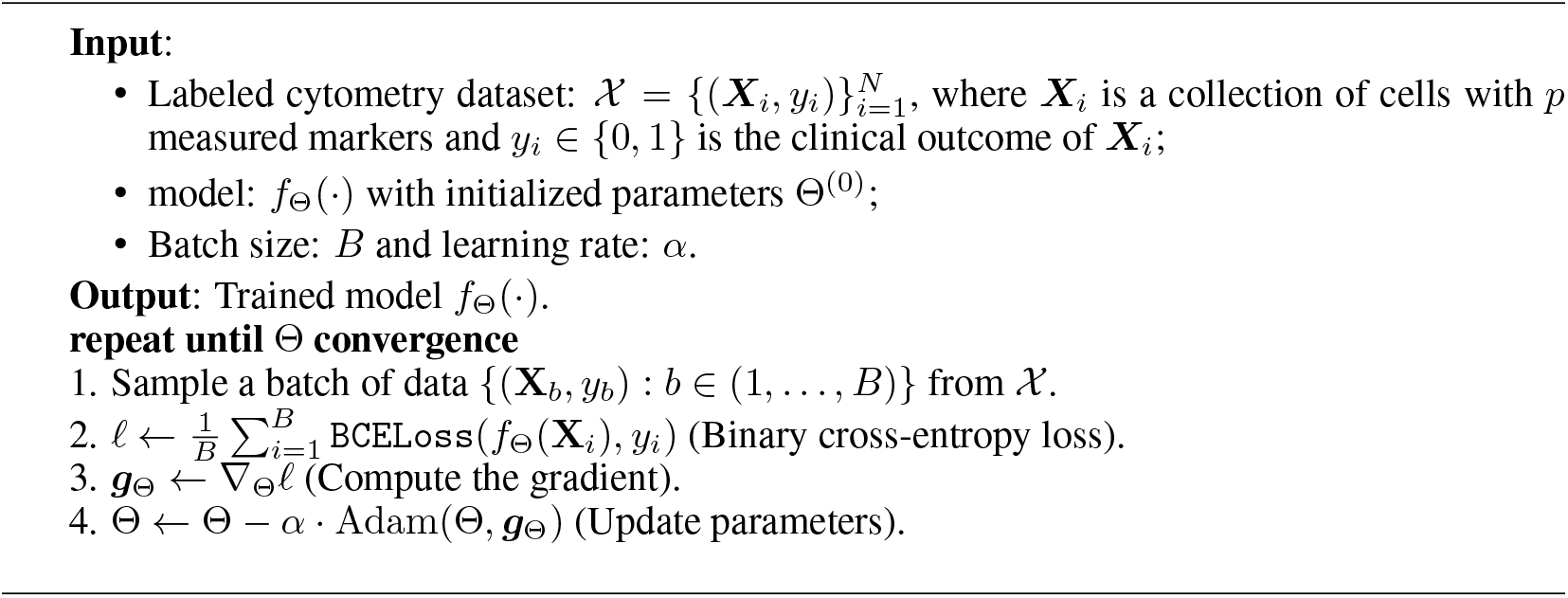
CytoSet Algorithm

### 2.6 Model Comparison

CytoSet and the two baseline methods, CellCNN (Arvaniti & Claassen, 2017) and CytoDx(Hu et al., 2019), all belong to the class of end-to-end gating-free methods. These methods all combine permutation equivalent and pooling functions to build a permutation invariant model on cytometry data. For the pooling function, CytoSet and CellCNN can either use max or mean pooling, while CytoDx only uses mean pooling. For the permutation equivalent function, both CellCNN and CytoDx use only one permutation equivalent layer while CytoSet can stack multiple layers. Thus, CellCNN and CytoDx can be viewed as special cases of CytoSet by limiting the model depth. The deeper architecture of CytoSet makes it more flexible for learning the complicated relationships between the raw input and the associated clinical outcome. This is because the model can reuse more information and increasingly generate abstract features from previous layers (Krizhevsky et al., 2012; He et al., 2016).

## 3 Results and Analysis

We ran a series of experiments to compare CytoSet to CellCNN and CytoDx on four flow and mass cytometry datasets. In section 3.1, we provide an introduction to the datasets used for the classification tasks. Next, in section 3.2, we provide details about the model training procedure, including the selection of the hyper-parameters and the model. Following in section 3.3, we evaluated our method and compare the classification performance with CellCNN and CytoDx. To ultimately make our model more interpretable, we visualized each sample according to their embedding vectors learned by CytoSet in section 3.5.

### 3.1 Description of Datasets

In our experiments, we compared the classification performance of CytoSet to two baseline methods (CellCNN (Arvaniti & Claassen, 2017) and CytoDx (Hu et al., 2019)) on four flow and mass cytometry datasets. Three of the tested datasets (HEUvsUE, AML, and HVTN) were used in the original FlowCAP-II benchmarking challenge (Aghaeepour et al., 2013).

#### HEUvsUE Dataset

The HEUvsUE dataset consists of 308 blood samples from African infants who were either exposed to HIV in *utero* but remain uninfected (HEU) or who were unexposed (UE). Results from the original FlowCAP-II challenge revealed that prediction in this dataset is quite challenging. In our experiments, we randomly selected 80% of the individuals for training and validation, and the remaining 20% for testing.

#### AML Dataset

The AML dataset has a total of 2872 samples collected from 359 AML (acute myeloid leukemia) and non-AML individuals. Similar to the experimental setup in the HEUvsUE dataset, 80% of the individuals were randomly selected for training and validation, and the remaining 20% were used for testing.

#### HVTN Dataset

The HVTN (HIV Vaccine Trials Network) dataset consists of 96 samples collected from one of two antigen stimulations (Gag and Env stimulated) in post-HIV vaccination T cells. In the experiments, we used the 50%-50% train-test split as indicated in the associated metadata in Ref. (Aghaeepour et al., 2013; Spearman et al., 2011).

#### NK-Cell Dataset

The NK-Cell Dataset (Horowitz et al., 2013) was previously used to evaluate CellCNN in Ref. Arvaniti & Claassen (2017). This dataset was pre-processed to include only cells across 20 samples that were characterized as NK cells, according to a biological expert. Moreover, the classification problem in this dataset is to classify Cytomegalovirus (CMV) seropositive and seronegative samples. In experiments, we adopted the same train-test split as the authors of CellCNN Arvaniti & Claassen (2017) to facilitate objective comparison.

#### Dataset Availability

The HEUvsUE, AML and HVTN datasets are all available through FlowRepository^1^ (Spidlen et al., 2012) with the following experiment IDs: FR-FCM-ZZZU (HEUvsUE), FR-FCM-ZZYA (AML), and FR-FCM-ZZZV (HVTN). The NK cell dataset is available with CellCNN (https://github.com/eiriniar/CellCnn). For more detailed statistics of these datasets, please refer to Table 5 in the Appendix.

### 3.2 Experimental Setup

#### Cytometry Data Pre-Processing

Before training, all of the raw measurements of cells were transformed using an `arcsinh` transformation (*f* (·)) with cofactor=5 (Azad et al., 2016). This transformation is defined as *f* (*x*) = arcsinh(*x*/5).

#### Evaluation Metrics

We used both the classification accuracy (ACC) and the area under the receiver operator curve (AUC) to quantify the classification performance.

#### Model Optimization

We trained our deep learning model using Adam optimizer (Kingma & Ba, 2014) with *β*_1_ = 0.9 and *β*_2_ = 0.999 across different datasets. The learning rate and batch size are set to 0.0001 and 200 across different experiments, respectively. In all of the experiments, we selected the best model based on the performance (in terms of AUC) on the validation dataset. To prevent overfitting, we also applied early stopping when training our models. As a result, the training stopped after five epochs of unimproved AUC on the validation dataset.

### 3.3 Results on Benchmark Flow and Mass Cyometry Datasets

#### 3.3.1 HVTN Flow Cytometry dataset

We trained our model on the HVTN dataset for 100 epochs using three permutation equivalent blocks. The test classification results and the corresponding ROC curves are reported in Table 1 and Figure 2(**a**), respectively. Our CytoSet model achieves the best performance in comparison to CellCNN and CytoDX. Further, both CytoSet and CellCNN outperform CytoDX by a large margin. In an experiment varying *k*, or the number of cells sampled from each patient’s set of cells, the performance across different models is only slightly changed.

**Table 1:** The testing ACC and AUC obtained from varying, *k*, the number of subsampled cells per patient sample in the HVTN dataset. The numbers reported are averaged over 5 runs with different random seeds.

| Model | $k = 256$ | | $k = 512$ | | $k = 1024$ | |
| --- | --- | --- | --- | --- | --- | --- |
|  | ACC | AUC | ACC | AUC | ACC | AUC |
| CellCNN (Arvaniti & Claassen, 2017) | 0.829 | 0.934 | 0.796 | 0.927 | 0.837 | 0.933 |
| CytoDx (Hu et al., 2019) | 0.658 | 0.746 | 0.629 | 0.738 | 0.654 | 0.741 |
| CytoSet | **0.913** | **0.963** | **0.892** | **0.964** | **0.883** | **0.957** |

**Figure 2:**
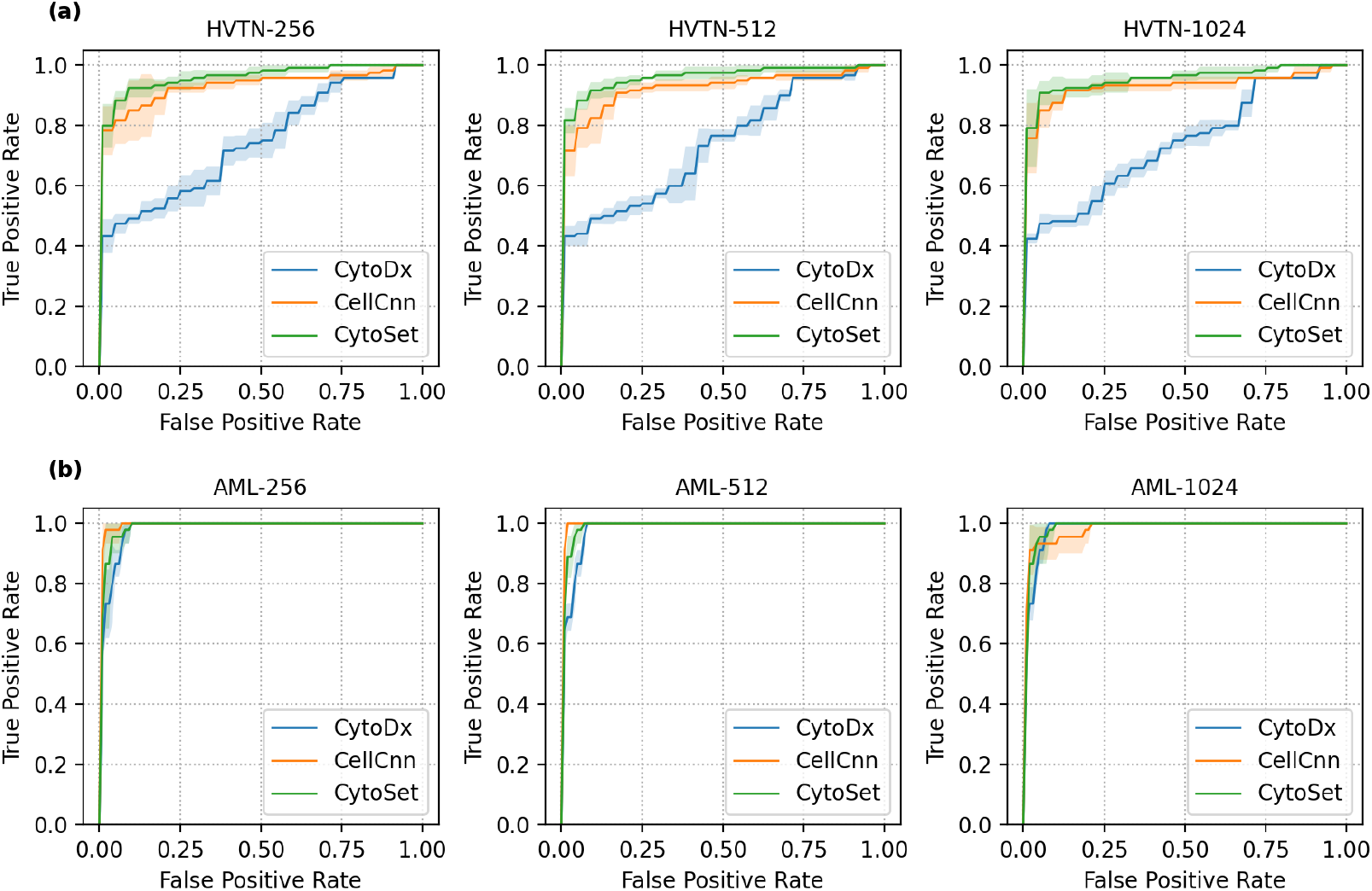
The test ROC curves of CytoDx, CellCNN and CytoSet on the HVTN (**a**) and AML (**b**) datasets from subsampling 256, 512, 1024 cells across patient samples, respectively. The shaded regions represent the standard deviation of 5 runs with different random seeds. All three methods perform comparably well on the AML dataset, but CytoSet outperforms CellCNN and CytoDX on the HVTN dataset.

#### 3.3.2 AML Mass Cytometry dataset

We trained our model on the AML dataset for 20 epochs using only one permutation equivalent block. We report the performance of the two baseline methods along with CyoSet in Table 2. The corresponding receiver operator curve (ROC) is illustrated in Figure 2(**b**). All of the methods perform reasonably well and exhibit comparable performance. This indicates that the AML dataset is relatively simple for models to fit. This point is also verified in the evaluation results of Ref. Aghaeepour et al. (2013). Again, the number of subsampled cells per patient sample, *k*, only slightly affects the classification performance of different models.

**Table 2:**
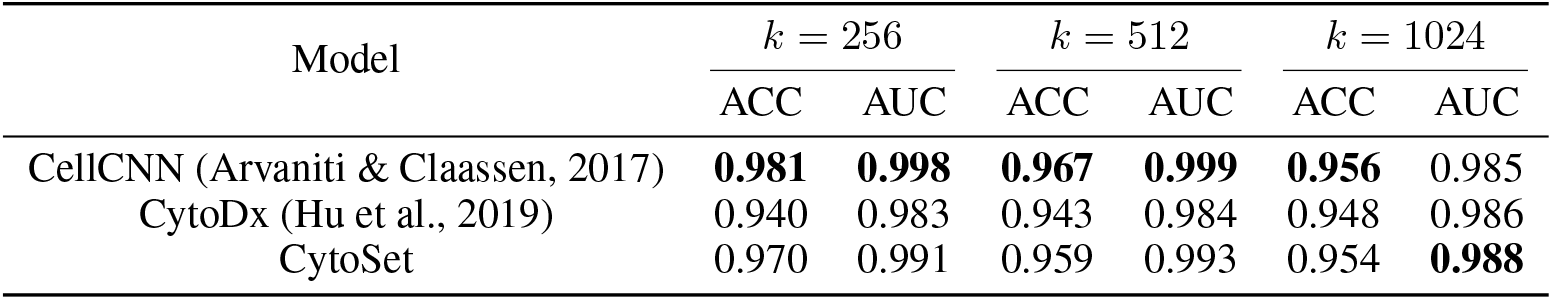
The testing ACC and AUC obtained from varying, *k*, the number of subsampled cells per patient sample in the AML dataset. The numbers reported are averaged over 5 runs with different random seeds.

| Model | $k = 256$ | | $k = 512$ | | $k = 1024$ | |
| --- | --- | --- | --- | --- | --- | --- |
|  | ACC | AUC | ACC | AUC | ACC | AUC |
| CellCNN (Arvaniti & Claassen, 2017) | **0.981** | **0.998** | **0.967** | **0.999** | **0.956** | 0.985 |
| CytoDx (Hu et al., 2019) | 0.940 | 0.983 | 0.943 | 0.984 | 0.948 | 0.986 |
| CytoSet | 0.970 | 0.991 | 0.959 | 0.993 | 0.954 | **0.988** |

#### 3.3.3 HEUvsUE Flow Cytometry dataset

We trained our model on the HEUvsUE dataset for 50 epochs using two permutation equivalent blocks. Considering the difficulty of fitting an appropriate model to this dataset, we used larger *k*s (e.g. the number of subsampled cells) in training. We report the test classification results in Table 3 and Figure 5 of the Appendix. CytoSet achieves the best accuracy (ACC) across different methods while CellCNN outperforms other methods in terms of AUC score. Varying *k*, or the number of cells sampled from each patient’s set of cells, the performance across different models is also robust.

**Table 3:** The testing ACC and AUC obtained from varying, *k*, the number of subsampled cells per patient sample in the HEUvsUE dataset. The numbers reported are averaged over 5 runs with different random seeds.

| Model | $k = 1024$ | | $k = 2048$ | | $k = 4096$ | |
| --- | --- | --- | --- | --- | --- | --- |
|  | ACC | AUC | ACC | AUC | ACC | AUC |
| CellCNN (Arvaniti & Claassen, 2017) | 0.565 | **0.790** | 0.565 | **0.754** | 0.638 | **0.754** |
| CytoDx (Hu et al., 2019) | 0.432 | 0.438 | 0.432 | 0.435 | 0.429 | 0.441 |
| CytoSet | **0.644** | 0.655 | **0.680** | 0.718 | **0.657** | 0.738 |

#### 3.3.4 NK-Cell Mass Cytometry dataset

In addition to the large datasets used in the FlowCap-II challenge, we also tested our model along with two baselines on the smaller NK-cell mass cytometry dataset. We trained our model for only 10 epochs using one permutation equivalent block, since there were only 14 training samples. The test classification results is shown in Table 4. Unlike other datasets, there is no consistent winner on this dataset. CytoSet and CytoDx outperform CellCNN on this dataset. The performance of CytoDx indicates that logistic regression is also good enough when there are only a few of samples available for training the classification model. The consistently strong performance of CytoSet across both large and small datasets shows that CytoSet generalizes well to diverse applications.

**Table 4:** The testing ACC and AUC obtained from varying, *k*, the number of subsampled cells per patient sample in the NK-cell mass cytometry dataset. The numbers reported are averaged over 5 runs with different random seeds.

| Model | $k = 256$ | | $k = 512$ | | $k = 1024$ | |
| --- | --- | --- | --- | --- | --- | --- |
|  | ACC | AUC | ACC | AUC | ACC | AUC |
| CellCNN (Arvaniti & Claassen, 2017) | **0.833** | 0.950 | 0.767 | 0.975 | 0.667 | 0.825 |
| CytoDx (Hu et al., 2019) | **0.833** | 0.925 | **0.833** | **1.000** | 0.833 | 0.850 |
| CytoSet | 0.800 | **0.975** | **0.833** | 0.975 | **0.867** | **0.950** |

### 3.4 Robustness Analysis

To verify the robustness of our model, we performed experiments varying the number of blocks in the models fit to the HVTN and HEUvsUE datasets. Here, we trained the models using 1, 2, and 3 permutation equivalent blocks (blue, orange, and green curves, respectively). The number of parameters used in different models is given in Table 6 of the Appendix. The test ROC curves in Figure 3 demonstrate that our CytoSet model is robust to the depth of the neural network.

**Figure 3:**
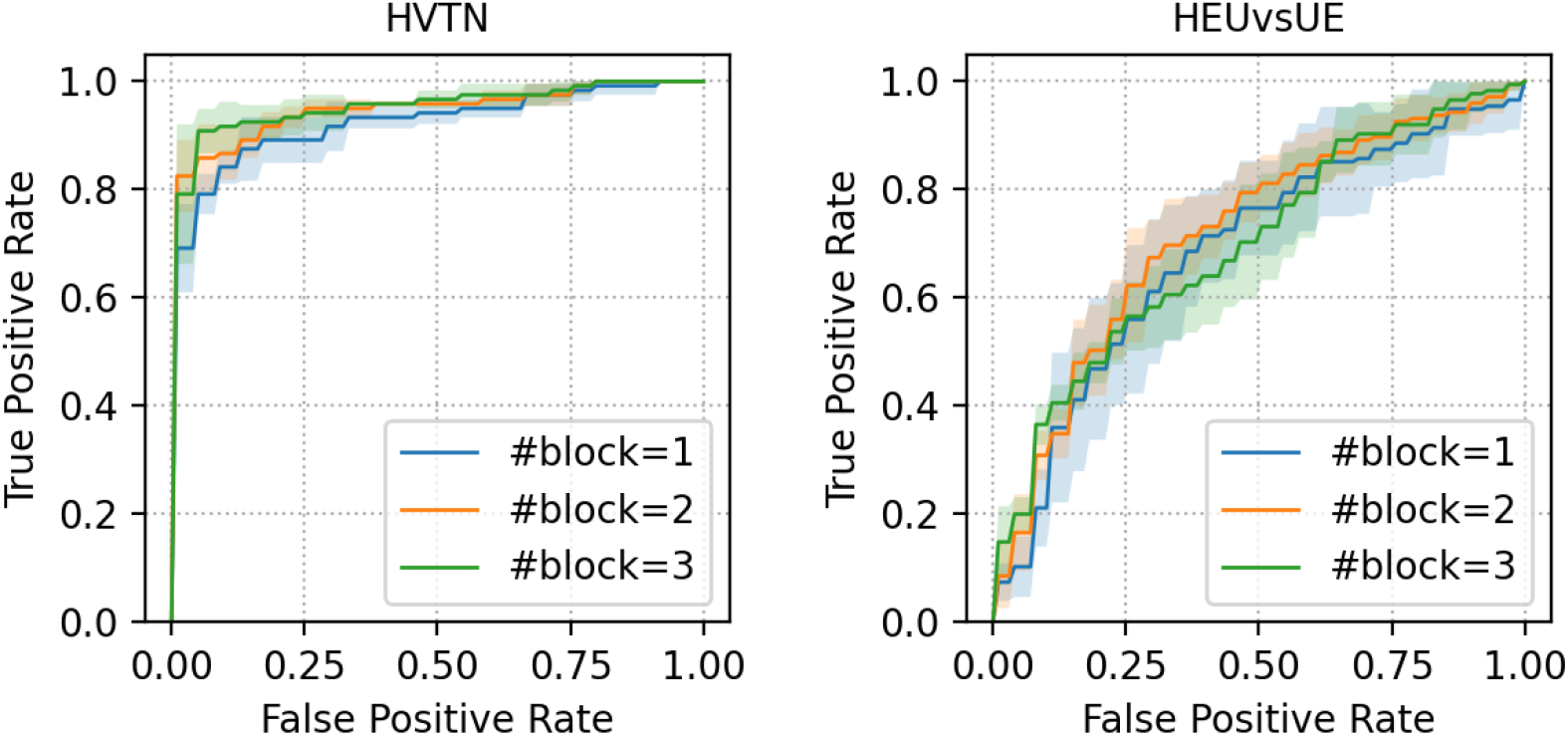
The test ROC curves of CytoSet applied to the HVTN and HEUvsUE datasets, trained using 1, 2, and 3 blocks (blue, orange, and green curves, respectively). The shaded region represents the standard deviation of five runs with different random seeds.

### 3.5 Visualization of Learned Sample Embeddings

To better understand the representations of each test samples learned by CytoSet, we visualized the set embedding vector, ***h*** in 2-dimensions with t-SNE (Van der Maaten & Hinton, 2008). As representative examples, we show visualizations for the AML and HVTN datasets in Figure 4. The t-SNE visualizations for the HEUvsUE and NK-cell datasets were omitted since the number of testing samples was quite small. In the AML dataset, the t-SNE visualization shows that the AML and healthy test samples tend to cluster together in 2-dimensional space. (Figure 4(**a**)). Similarly, in the HVTN dataset, GAG-stimulated and ENV-stimulated test samples also cluster by class (Figure 4(**b**)). These observations qualitatively demonstrate that CytoSet can learn the effective representations of the sets for discriminating the associated clinical outcomes.

**Figure 4:**
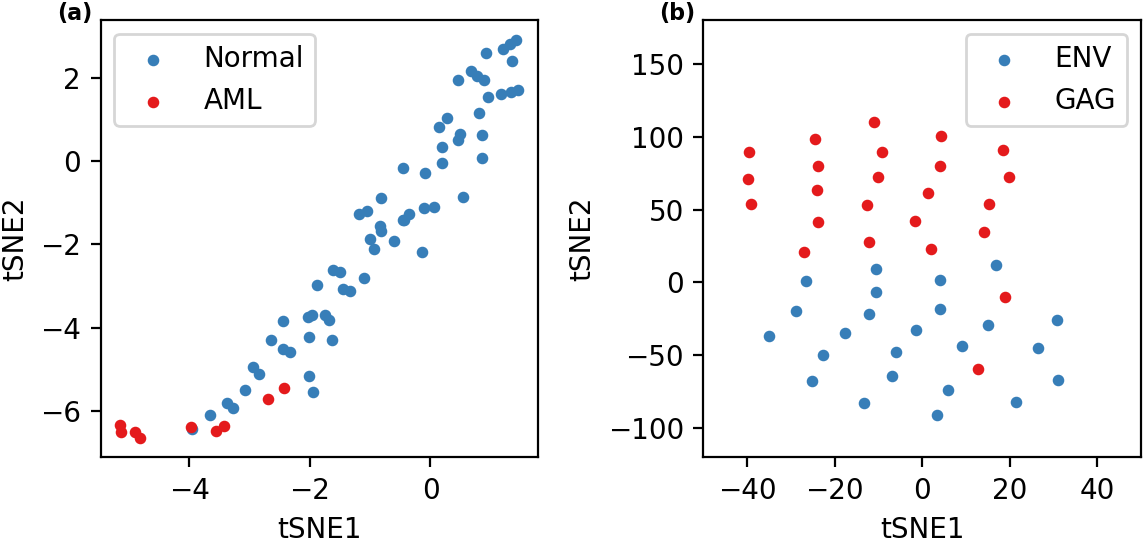
t-SNE visualizations of the samples from patients in test set in the AML (a) and HVTN (b) datasets. Samples were projected into two dimensions by applying tSNE to the set embedding vector, ***h***, learned by CytoSet. The color of each point (patient sample) indicates the phenotype (clinical outcome).

## 4 Conclusion

In this paper, we proposed a new deep learning model called CytoSet for predicting clinical outcomes from single-cell flow and mass cytometry data. In the problem setup, CytoSet innovatively formulates the prediction of clinical outcomes as a classification task on sets. For the architecture design, CytoSet generalizes two gating-free methods CellCNN and CytoDx using stackable permutation invariant network architectures, which improves the model’s expressive power.

In the experiments, we demonstrate that CytoSet outperforms two other gating-free methods on the classification task of a large cytometry datasets, HVTN and is also comparable to CellCNN on other benchmark datasets. We also show that CytoSet is robust to the *k*, or the number of subsampled cells and the network depth while training the model on large and challenging datasets such as HVTN and HEUvsUE. CytoSet successfully advances the ability to represent and link data generated through single-cell assays, such as flow and mass cytometry, to external clinical variables of interest.

## Acknowledgments

We would like to thank the ITS at The University of North Carolina at Chapel Hill for providing computational resources to run the experiments in this paper. We also gratefully acknowledge the input from the Reviewers who provided us with valuable feedback.

## A Appendix

**Table 5:** The details of the datasets used in the experiments.

| Scale | Name | Cytometry | #Samples | #Patients | Train/Test Split |
| --- | --- | --- | --- | --- | --- |
| Large | AML | Mass | 2872 | 359 | 4:1 |
| Large | HEUvsUE | Flow | 308 | 44 | 4:1 |
| Medium | HVTN | Flow | 96 | 96 | 1:1 |
| Small | NK Cell | Mass | 20 | 20 | 7:3 |

**Figure 5:**
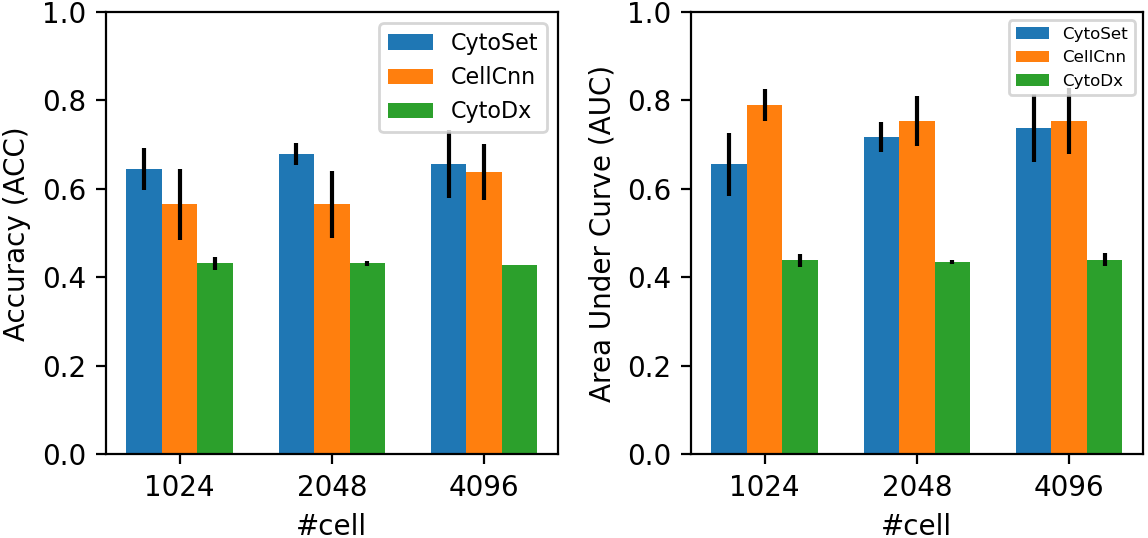
The accuracy (left) and area under curve (right) of CellCNN, CytoDx and CytoSet on the HEUvsUE dataset from subsampling {1024, 2048, 4096} cells across patient samples, respectively.

**Table 6:** The number of parameters of the CytoSet model on the HEUvsUE dataset with various numbers of blocks.

|  | #block=1 | #block=2 | #block=3 |
| --- | --- | --- | --- |
| #Params($\times 10^3$) | 71.4 | 202.8 | 334.1 |

## Footnotes

1 https://flowrepository.org

